# Focused Ultrasound Crosslinkable Granular Hydrogels

**DOI:** 10.1101/2025.04.14.648397

**Authors:** Natasha L. Claxton, Estelle He, Kelly Bukovic, Eric A. Thim, Rachel A. Letteri, Steven R. Caliari, Richard J. Price, Matthew R. DeWitt, Christopher B. Highley

## Abstract

Minimally invasive delivery of biomaterials for tissue regeneration can be achieved using biomaterials delivered by injection that rapidly stabilize at the delivery site. Tissue regeneration has been shown to be dependent on material properties, with porosity strongly supporting revascularization and tissue regrowth. Here, a highly porous granular hydrogel system was designed that is compatible with the unique strengths of highly penetrating, minimal invasive focused ultrasound (FUS), to address this challenge. FUS offers well-defined spatial and temporal control over hydrogel crosslinking. We developed a composite granular hydrogel scaffold composed of two polyethylene glycol (PEG)-based components, microgels and fibers with FUS-responsive chemistry, and a pore-defining gelatin microgel component. Upon applying FUS, the bulk granular hydrogel stabilized through the formation of crosslinks between PEG components, with porosity designed by gelatin microgel melting. FUS-crosslinking parameters were determined that resulted in crosslinking both *in vitro* and a mouse cadaver model of minimally invasive delivery. The resulting granular hydrogels’ stability depended on the presence of fibers and exhibited viscoelastic properties comparable to granular hydrogels that were photocrosslinked. Hydrogels were highly porous and cytocompatible. This work defines a FUS-responsive granular system and extends the potential of FUS as a novel, noninvasive method for crosslinking regenerative hydrogel systems.

## 1. Introduction

Hydrogels are well established in regenerative medicine due to their properties that closely mimic natural tissues and potential for application-specific design. Their high water content and a three-dimensional (3D) network structure is similar to the extracellular matrix (ECM) in human tissues. Additionally, their mechanical properties and degradation rates can be precisely engineered to match target tissue and regeneration. Hydrogels have been used in diverse applications across tissue systems in wound healing, cartilage repair, bone regeneration, addressing cardiac tissue injuries, and to support neural tissue regeneration. In many medical contexts, minimally invasive delivery of a hydrogel by injection is desirable, which requires that hydrogels transition rapidly from a liquid-like to gel state *in situ* after leaving the needle or catheter used in delivery. To enable the delivery of hydrogel precursor solution and the formation of a stable nanoporous network at the site of injection, rapid hydrogel crosslinking is required. Hydrogels have been engineered to crosslink through mechanisms such as matching crosslinking kinetics to timing of delivery, mixing during delivery, pH changes, electrostatic interactions, or enzymatic reactions^1–4^. Nonetheless, achieving precise gelation at a delivery site remains a significant challenge due to requirements of spatiotemporal control of crosslinking. Externally triggered crosslinking by techniques such as photoinitiated reactions provides some access to controlling the onset of crosslinking, but in minimally invasive applications, external triggers might not sufficiently penetrate tissue to a delivery site ^5,6^, leaving the issue unresolved.

Ultrasound technologies offer a simple, direct, controllable, non-invasive stimulus^7,8^ that can be used to alter structural characteristics of hydrogels^9^ and trigger crosslinking. Ultrasound has the capability to polymerize hydrogel formulations^10–15^ and control porosity of engineered scaffolds^15–17^. It can also initiate dynamic changes in thermally responsive biomaterials^14,18^ that change conformation based on temperature resulting globule form in heated environments and back to coiled conformation upon cooling. Ultrasound energy penetrates deeply in tissue, and thus is promising in minimally invasive application. Recently, focused ultrasound (FUS), which directs ultrasonic waves to a focal point with millimeter-scale resolution, has emerged as a means of externally triggering minimally invasive hydrogel crosslinking with spatial and temporal control^19,20^. FUS leverages both thermal and mechanical effects, enabling deep tissue penetration (>100 times deeper than light into optically scattering materials)^21^ without affecting, or heating, tissue along the beam path and remaining noninvasive and nonionizing^10,20^. FUS can effectively generate reactive oxygen species (ROS) ^13,21,22^, which can facilitate hydrogel formation within minutes^21^.

In recent hydrogel applications, FUS has been used for drug and protein delivery by selectively tuning hydrogel chemistry reactions to control the release rate and timing without degradation kinetics^12,23,24^. When applied to biomaterials, FUS sonications can induce reactions that result in material functionalization^11,14,16,25^. Additionally, FUS has been used in a 3D-printing type process, where targeting of the US focal point in 3D space can spatially control polymerization of hydrogel precursor solutions, or sono-inks^21^, present at the focal point, yielding complex structures and overcoming depth limitations related to light-activated in 3D printing in similar applications. While the versatility of FUS has been demonstrated, using it in conjunction with injected hydrogel precursor solutions may still face challenges in certain applications. These solutions remain flowable when injected and may flow from the site of delivery prior to stabilization by FUS, which can take minutes. Upon crosslinking, hydrogel precursor solutions also typically form structures with nanoscale porosity, which may require additional design for degradation to support tissue regrowth and vascularization.

Granular hydrogel systems present an opportunity to address these limitations, as they have been established in applications combining injectability with the ability to form scaffolds that maintain stable pore structures essential for tissue integration and nutrient support. Granular hydrogels, whose macroscale bulk is formed from discrete hydrogel particles (or microgels), can flow during injection and stabilize upon delivery by physical forces among particles as well as interparticle crosslinking, or annealing. In seminal work using these materials in regenerative medicine and wound-healing applications, microporous annealed particle (MAP) hydrogels enhanced tissue regrowth^9^ and have demonstrated favorable immunological responses^26^. Recently, the inclusion of electrospun hydrogel fibers has been shown to provide additional stability in annealed granular hydrogels, including enabling mesoscale (100 µm – 1 mm) pores to be sustained within these materials^27^. This pore content can be defined by design in a material that is also injectable by the inclusion of a removable gelatin microgel. Ratiometric control of the proportion of removable microgels to the non-removable microgel and fiber component allows large, interconnected pores to be formed after injection. The use of a gelatin microgel component avoids compaction of pore spaces, which can occur when porosity is increased in granular systems with the dilution of microgels. This issue arises during extrusion or injection, where dilute systems can compact with interstitial fluid used for dilution, causing separation from the hydrogel^28^. Long range particle-fiber interactions can stabilize mesoscale pores over time and allowing design of scaffolds that support high cellularity^27^.

In this work, we show that FUS can be used to crosslink granular hydrogel materials *in situ* using a fiber reinforced scaffold, and we show that the resulting materials are highly porous, toward ultimately supporting wound healing and tissue regeneration. Specifically, we utilized FUS to trigger crosslinking through excess acrylate and norbornenes functional groups present on granular materials. Our composite granular materials included polyethylene glycol (PEG)-based microgels with excess acrylate groups, PEG-based electrospun fibers with excess norbornene groups, sacrificial gelatin microgels, and thermoresponsive polymers. The thermal effects of FUS melted the gelatin microgels and FUS mechanisms induced crosslinking at the surfaces of microgels and fibers *in vitro*. To evaluate this system’s potential for use *in vivo*, we injected the hydrogel formulation subcutaneously into a mice cadaver model and observed that FUS induced crosslinking and generated stable fiber-reinforced hydrogels localized to subcutaneous injection sites. This approach highlights the potential of FUS to be used in conjunction with granular hydrogels for translational applications and therapeutic purposes.

## 2. Results and Discussion

### 2.1 Designing FUS-Responsive Granular Hydrogels

The granular hydrogels used in this work were composed of PEG microgels with excess acrylates, PEG fibers with excess norbornenes, and gelatin microgels (**Figure 1A**). In this system, PEG microgels and fibers were designed to compromise the final structure of granular hydrogels, with gelatin microgels intended to be a removable component. The inclusion of the gelatin microgels to create porosity, as opposed to simply diluting the PEG components, allows design control over pore fraction in the final gel. Removable microgels also avoid a loss of porosity during injection processes that can occur in dilute granular hydrogel formulations. Soluble PEG-thiol was included to form crosslinks between PEG microgels and fibers within granular hydrogel scaffold upon FUS sonication. Based on previous work^11^, we hypothesized that FUS sonication, in the presence of a free radical initiator (potassium persulfate, KPS) that generates radicals in response to both light-and thermal-based cues, and a thermally responsive acoustic absorber material^27^ that reduces convective motion upon heating (poly(di(ethylene glycol) methyl ether methacrylate, PDEGMA), would support interparticle crosslinking at the focal point of ultrasonic waves (**Figure 1B**). Radicals generated by FUS radicals ^29^ can initiate thiol-ene chemical reactions near the focal point, forming crosslinks between the microgel and fiber components through the soluble PEG-thiol molecules (**Figure 1B, II**). PDEGMA undergoes a conformational structural change with increasing temperature and was expected to respond to FUS-induced heating, resulting in increased viscosity. By locally increasing viscosity to slow molecular motion^21^ and convective flows, PDEGMA would support crosslinking reactions (**Figure 1A-B, II**). Also, upon applying FUS, gelatin microgels were expected to liquefy (melting point: 37°C) as crosslinking occurred, due to the combination of temperature and radical generation at the FUS focal point (**Figure 1B**). After FUS application, a crosslinked granular hydrogel will remain with pores formed by gelatin microgels melting (**Figure 1C**). After sonication, soluble gelatin and PDEGMA, which returns to its original conformation are expected to remain in solution and gradually diffuse away from the stabilized granular hydrogel over time.

**Figure 1.**
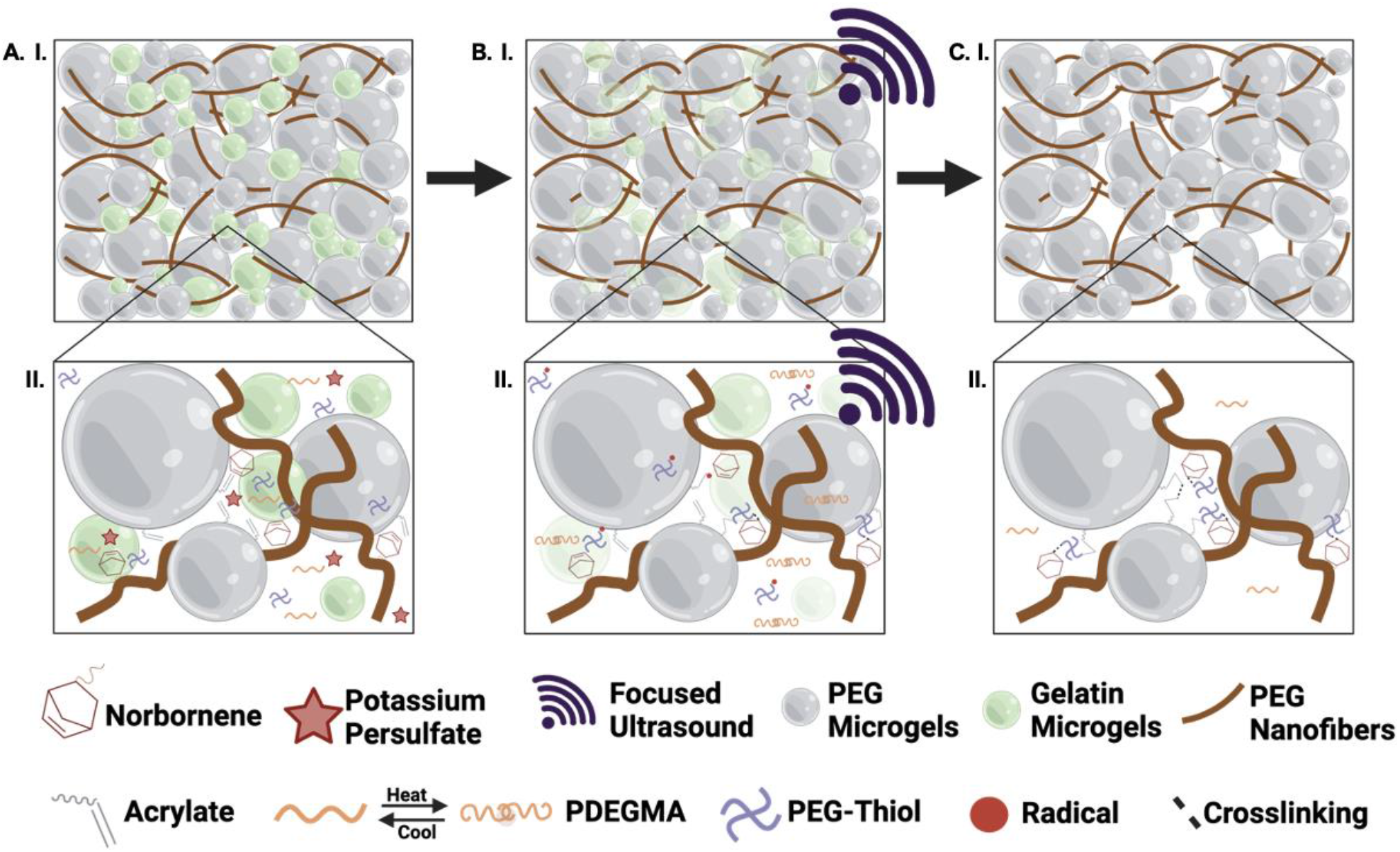
Schematic illustration of the focused ultrasound crosslinking mechanism for granular hydrogel scaffolds. **A**. Polyethylene glycol (PEG) microgels containing excess acrylates, PEG fibers containing excess norbornenes, and gelatin microgels (***I***) are mixed with a PEG-thiol solution, an initiator (potassium persulfate), and a thermo-responsive material (PDEGMA) (***II***). **B**. Upon focused ultrasound sonications, gelatin microgels begin to melt (***I***), radicals form on the monomers initiating crosslinking, and PDEGMA undergoes globule structural transformation upon heating (***II***). **C**. After focused ultrasound stimulus, gelatin microgels fully melt leaving behind PEG microgels and fibers (***I***) that are completely crosslinked resembling a continuous, bulk hydrogel. PDGEMA goes back to its original structure upon cooling (***II***).

### 2.2. Crosslinking PEG Hydrogel Precursors and PEG Granular Hydrogels with FUS

To confirm the FUS parameters for triggering reactions that would drive interparticle crosslinking, we first evaluated crosslinking of a continuous, bulk hydrogel before moving to a granular formulation, where a lack of stabilization via annealing crosslinking might not reflect non-reactivity. Continuous PEG hydrogels have been crosslinked using ultrasound^30,31^, and building on published results^10,11^, we prepared a hydrogel precursor solution containing 20 wt% PEG-diacrylate (PEGDA), KPS as a radical generator, and PDEGMA as a thermoresponsive acoustic absorber. We sonicated samples in microcentrifuge tubes with FUS, and continuous hydrogels were visually observed within 10 minutes. As a measure of gelation, the resulting hydrogels were measured to have storage moduli of 10 kPa. To confirm crosslinking via FUS was similar to established photocrosslinking, we compared hydrogels formed via FUS to hydrogels formed from the same precursor solution using photocrosslinking. Photocrosslinked gels had similar stiffnesses of 11 kPa (**Figure 2A, I**), providing evidence that the FUS conditions could drive the formation of crosslinks within hydrogels networks where resulting hydrogels had properties analogous to those achieved by photocrosslinking.

**Figure 2.**
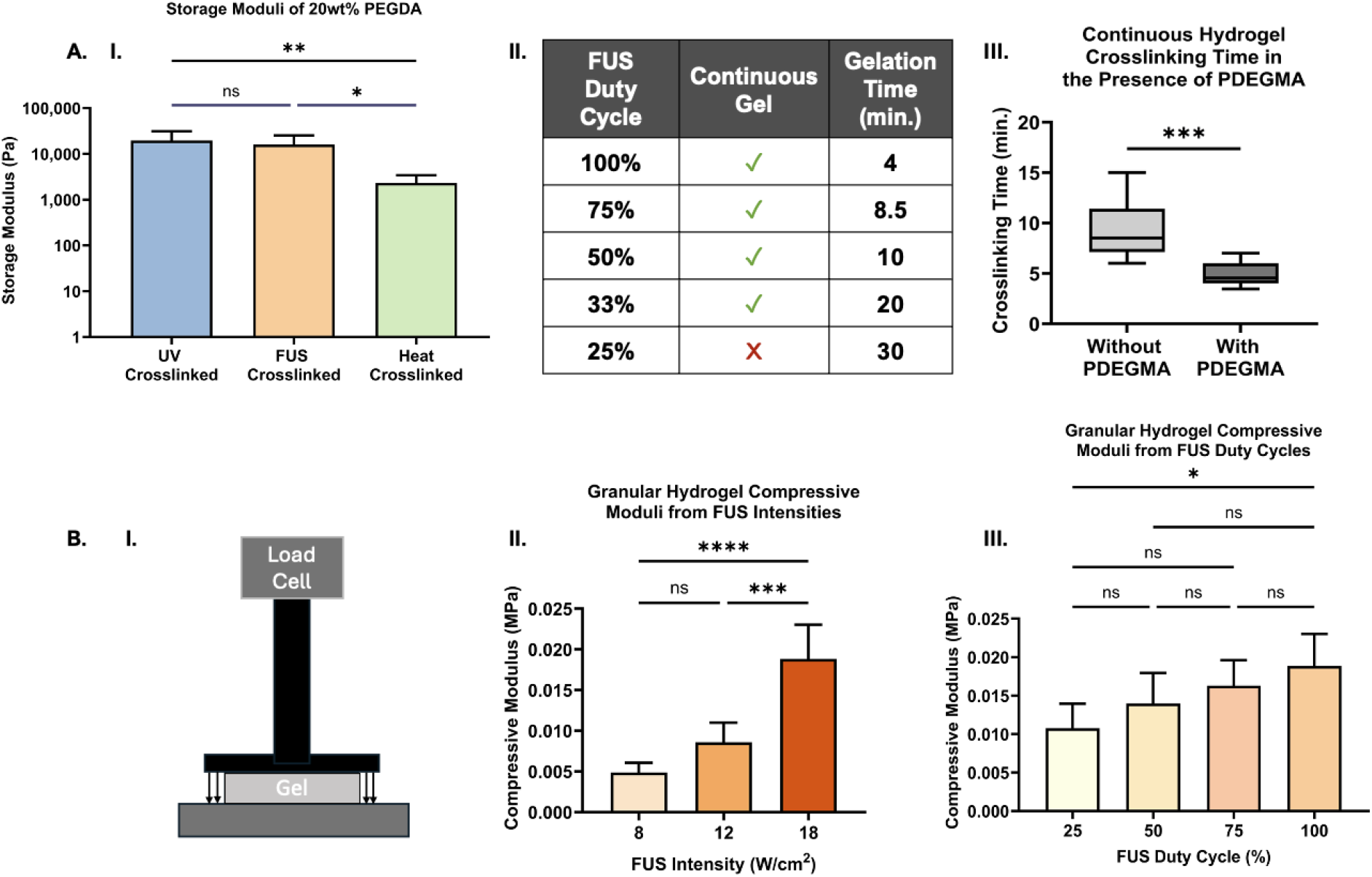
Focused ultrasound responsiveness on precursor solution and granular hydrogel materials. **A**. An oscillatory time sweep of 20 wt% PEG-diacrylate hydrogels crosslinked via UV light shows similar stiffness compared to the FUS crosslinked gel. Heat crosslinked gel exhibits a softer gel compared to UV and FUS crosslinked gels (***I***; One-way ANOVA with Tukey’s posthoc test, * P<0.05, **P<0.005). Continuous hydrogel crosslinking with FUS was achieved with duty cycles as low as 33% (***II***). In the presence of a thermoresponsive material, PDEGMA, crosslinking time was faster than without PDGEMA (***III***; T-test, *** P < 0.0001). **B**. Determination of compressive moduli using Instron compression testing using quasi-static compression testing of cylindrical pucks of crosslinked granular hydrogel material (***I***). Crosslinking of granular hydrogel achieved from varied FUS intensities (***II***) and the degree of compressive stiffness increases with varied FUS duty cycles

Next, we wanted to determine whether the crosslinking observed was due to KPS’ thermal responsiveness to heat generated by FUS alone or to what extent the response of the aqueous environment to FUS initiated crosslinking through KPS by other physical mechanisms related to FUS, such as free radicals and photons local to collapsing cavitation bubbles^32^. To test this, we used KPS to crosslink our hydrogel precursor thermally. The resulting PEGDA gel exhibited a stiffness of 1.5 kPa (**Figure 2A, I**). The significant decrease in mechanics when using only a thermal stimulus similar to the heating resulting from FUS indicates a substantial contribution of non-thermal energy to the crosslinking observed when using FUS to trigger radical generation and acrylate polymerization in the precursor solution.

Next, we wanted to further understand how FUS parameters might be controlled to generate crosslinking and were specifically interested in the extent to which duty cycle might be modulated to potentially reduce localized healing while still crosslinking the precursor solution. The duty cycle is the percent of time ultrasound waves are actively being transmitted from the transducer, and lower duty cycles result in decreased local heating. We applied FUS waveforms including a continuous (100%) wave and pulsed FUS waveforms corresponding to 75%, 50%, 33%, and 25% duty cycles. We obtained a continuous, bulk PEGDA hydrogel with duty cycles as low as 33% duty cycle, but gelation was not seen with as 25% duty cycle (**Figure 2A, II**). Decreased duty cycles corresponded to longer times observed for hydrogel gelation. To test the importance of having an acoustic absorber, PDGEMA, in the system, we compared FUS crosslinking of continuous hydrogel precursor solutions with and without PDEGMA. We observed that it took 2-3x longer (10-12 minutes) to crosslink a continuous PEGDA hydrogel in the absence of PDEGMA (**Figure 2A, III**).

We next considered granular hydrogels, formulated as described, whose crosslinking we confirmed through mechanical analysis. After FUS sonications, samples underwent compressive mechanical testing via Instron (**Figure 2B, I**). To evaluate material stiffness differences in the granular hydrogel crosslinking and determine the minimal intensity required for crosslinking, FUS intensity was modulated while maintaining a 100% continuous FUS duty cycle. At the lowest input voltage of 8 W/cm^2^, a stable gel with compressive modulus of 5 kPa gel was obtained compared to 12 W/cm^2^ and 18 W/cm^2^ yielding granular hydrogels with moduli of 10 kPa and 20 kPa, respectively (**Figure 2.B, II**). These data indicate that increasing FUS intensities are necessary to induce more complete crosslinking, resulting in stiffer granular hydrogels.

By varying FUS duty cycles, we observed increased compressive moduli for the crosslinked granular hydrogel as FUS duty cycle increased. Within a granular hydrogel system, a 25% duty cycle yielded a stable granular hydrogel with a bulk stiffness of 10 kPa compared to the continuous, 100% duty cycle FUS waveform that resulted in granular hydrogels with 19 kPa stiffnesses (**Figure 2.B, III**). The results indicate a modest trend in increasing moduli with duty cycle, suggesting increasing interparticle crosslinking within the system as duty cycle increased. However, a statistically significant difference is only observed between the 25% and 100% duty cycle groups. Because duty cycle decreases result in lower temperatures^33^, these data also suggest that the contribution of thermal energy to crosslinking is likely modest and not necessary for crosslinking to occur. A stable hydrogel with ∼12 kPa compressive moduli in the absence of significant thermal deposition in the 50% duty cycle group. This suggests that the crosslinking of the granular material is primarily due to ROS production through interactions of KPS with the FUS energy. While previous methods have demonstrated the ability to crosslink hydrogels using focused ultrasound^33^, they often rely on high power levels that generate substantial heat to induce crosslinking. While useful for 3D printing applications, this approach poses a risk *in vivo*, especially near thermally sensitive structures such as vasculature and nerves, making our low-power approach particularly advantageous for safer and more controlled in situ crosslinking. Future optimization of duty cycle and ultrasound intensity will enable crosslinking of gel *in vivo* without significant risk to collateral tissue.

### 2.3 FUS-Crosslinked, Fiber-enhanced Granular Hydrogels Porosity and Stability

Having confirmed the formation of mechanically robust fiber-reinforced granular hydrogels with FUS-induced crosslinking, we next considered hydrogels’ porous microstructure, the effects of the fiber component on stability, and cellular integration into pore space *in vitro*. It has been shown that ratiometric incorporation of gelatin microgels among the particle and fiber components allows the fraction of the pore space to be designed^27^, and for the large continuous pores to form upon melting gelatin. To confirm that FUS directly melts gelatin microgels, we tested the effects of FUS on gelatin microgels containing a black dye (for visualization) in a microcentrifuge tube. Before FUS was applied, these gelatin microgels demonstrated solid-like properties as a packed, granular hydrogel with a microcentrifuge tube (**Supporting Video 1**), the granular system did not exhibit flow during handling. Immediately after FUS stimuli, the granular hydrogel was observed to have melted, becoming a continuous, flowing fluid (**Supporting Video 2**).

When a fiber-reinforced granular scaffold is formed with gelatin microgels as a component of the system, it has been shown that the fibers are necessary and sufficient to sustain the highly porous microstructure that emerges in the granular system when gelatin melts. To visualize the hydrogels’ porosity, we imaged FUS-crosslinked and photocrosslinked gels by scanning electron microscopy (SEM) and confocal imaging. Scaffolds were crosslinked via UV light and FUS sonications. Samples placed under UV light were incubated at 37°C to melt gelatin microgels. SEM images show that the FUS crosslinked gel exhibits larger pores compared to UV crosslinked gel (**Figure 3A**). This is likely due to the movement of material as FUS is being applied. Additionally, the longer times to crosslink scaffolds using FUS compared to using light suggest that the physics of radical generation using FUS result in slower net crosslinking, with the rate of total bond formation being lower. This could result in weaker overall scaffolds during the crosslinking time course compared to processes using photoactivation of the radical initiator. Combined with FUS induced flows, material movement during the process might promote multivalent interactions to stabilize surface interactions, which could be seen in increased fiber and microparticle accumulation around pores.

**Figure 3.**
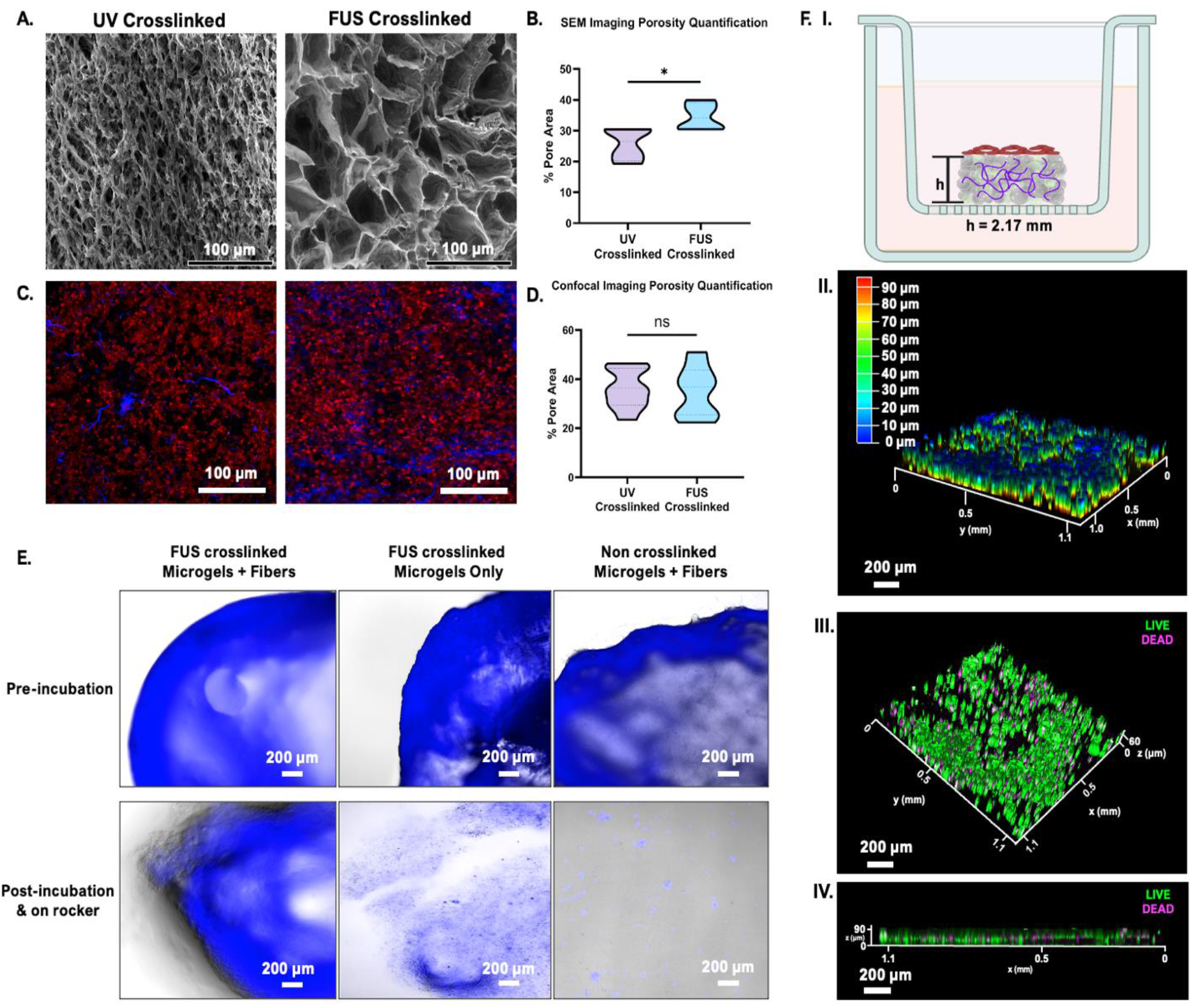
Material properties of granular hydrogel system post-FUS. **A**. SEM images of pore spaces within UV crosslinked and FUS crosslinked granular hydrogels. (scalebars = 100 µm). **B**. FUS crosslinked granular hydrogels yield higher percentages of pore area within the scaffold compared to the UV crosslinked scaffold. **C**. Confocal images of granular hydrogel pore space of UV crosslinked and FUS crosslinked scaffolds (blue – fibers and red – microgels). Images taken at 10x magnification. (scalebars = 100 µm). **D**. Quantification of pore space in confocal images. **E**. Images of granular hydrogel scaffolds pre- and post-incubation of FUS crosslinked and non-crosslinked scaffolds after being on rocker for 12 hours. FUS crosslinked granular materials maintain shape and structure after incubation and being on the rocker compared to microgels only and non-crosslinked granular material where it falls apart. Images show the importance of fibers for scaffold structure integrity. (scalebars = 200 µm). **F**. Schematic of HUVECs seeded on top of a FUS crosslinked granular hydrogel with a height of 2.17 mm in a transwell (***I***). Depth of HUVECs that travelled through the scaffold after 2 days are shown with majority of the cells at the bottom of the transwell (***II***). Angled (***III***) and side view (***IV***) images of HUVECs in 3D FUS crosslinked granular hydrogel via live/dead staining (Live – green, Dead – magenta) after 2 days post seeding. (scalebars = 200 µm).

Quantification of porosity in SEM micrographs shows that pore spaces within the FUS crosslinked granular hydrogels contribute to ∼30-40% of the total area seen, compared to 20-30% in the photocrosslinked materials (**Figure 3B**). To visualize pore space via confocal imaging, we formulated PEG microgels fluorescently tagged with thiolated rhodamine-B and PEG fibers tagged with 5(6)-carboxyfluorescein amidite (FAM). Again, samples crosslinked by FUS were compared to photocrosslinked samples. Microgels (red) and fibers (pseudocolored blue) distributions appear similar (**Figure 3C**). Quantification of unlabeled pore spaces in confocal images show similar porosity area coverages (**Figure 3D**). Both methods of crosslinking exhibit similar porosity area coverages (**Figure 3D**). This suggests that when scaffolds remain continuously hydrated, their material architectures remain similar independent of the crosslinking mechanism. The slight increase of pore space, as quantified using SEM, may be due to the effects of ultrasonic treatment, which, at higher powers (> 40W) than used here, has been shown to cause increases in the mean pore size, pore interconnectivity, and porosity in tissue scaffolds^34^.

To further illustrate the importance of both FUS crosslinking and the inclusion of fibers to the stability of these materials, we compared granular hydrogels’ stability against erosion of microgel components into surrounding media in three different groups: a FUS-crosslinked granular hydrogel containing microgels and fibers, a FUS-crosslinked granular hydrogel containing only microgels (no fibers), and a granular hydrogel containing microgels and fibers that was not FUS-crosslinked. Each resulting hydrogel was incubated on a dish on a rocker for 12 hours, and imaged before and after (**Figure 3E**). The FUS-crosslinked system containing fibers was seen to remain largely intact after 12 hours. The FUS-crosslinked system that did not contain fibers evidenced significant erosion. The hydrogels containing fibers and particles that were not exposed to FUS were not intact. Based on these differences, the fibers were not necessary just for stabilizing scaffold porosity in the granular system, but also for allowing FUS sonication to be used to crosslink the particle-based system studied here.

Toward supporting tissue regeneration, we assessed *in vitro* whether cells can move into the pore spaces of a FUS crosslinked granular hydrogel using the same ratio of gelatin (75:25 PEG:gelatin microgels) for non-cellular FUS experiments. We seeded HUVECs on top of a FUS-crosslinked granular hydrogel with a 2.17 mm height (**Figure 3F, I**) and observed cellular viability and locations in the scaffold after 48 hours. The majority of endothelial cells within the bottom ∼90 µm of the scaffold (denoted by red color in depth graph) (**Figure 3F, II**), indicating they were easily able to traverse the highly porous system. Viability staining for living and dead cells indicated that a majority of the HUVECs survived (**Figure 3F, III**) and were evenly distributed throughout the bottom of the granular hydrogel scaffold (**Figure 3F, IV**). To confirm cells were moving through the scaffold and not flowing around and infiltrating from the bottom, we applied cells in a small volume of medium to the top of the scaffold and observed cell distribution 15 minutes after seeding. Cells were found at the bottom of the scaffold after only 15 minutes, suggesting that large pores sizes were large enough to support convection flow of cell suspended in medium through the material. This degree of porosity should ultimately be supportive of cell migration into scaffolds and tissue growth within the pore spaces when considering *in vivo* applications. The porosity achieved in this scaffold via gelatin microgels show potential promise for tissue and microvasculature ingrowth *in vivo* similar to granular hydrogel *in vivo* work already conducted^9,26,35,36^.

### 2.4 Drug Release from Granular Hydrogels using FUS Sonications

This system has the potential to be used in minimally invasive therapeutic treatments in regenerative medicine applications through the combination with FUS. To enhance its therapeutic effects, we explored possibilities for drug release from the granular hydrogel system to support disease applications or regenerative medicine applications where efficacy might be improved by delivering soluble compounds in combination with FUS^37^. A model drug (doxorubicin, DOX) was incorporated into both the PEG and gelatin microgel components. We expected FUS could thus be used to induce rapid release from the gelatin particles through melting, with continued release coming from the PEG microgel component within the stabilized system. When quantifying DOX release, we observed ∼20% of DOX was released from the granular hydrogel scaffold (**Figure 4A**) immediately after FUS sonications within the first 6 hours (**Figure 4A**, inset). In samples containing gelatin microgels only (**Figure 4B**), we observed melting and 100% release after FUS exposure. While some burst release from the PEG microgels contributes to the burst seen for the combined granular hydrogel system release from the PEG microgel component continues for days afterwards (**Figure 4A**). From the entire granular hydrogel system, we observe that the gelatin microgels alone released ∼2.5% of the theoretical DOX encapsulated, while the PEG microgels released ∼16% of the theoretical DOX encapsulated after 48 hours (**Figure 4B**). The remainder of the theoretical DOX encapsulation in the gelatin microgels was lost in washing steps during processing. It is expected that a fraction in excess of 80% was also lost during processing the PEG microgels due to washing steps.

**Figure 4.**
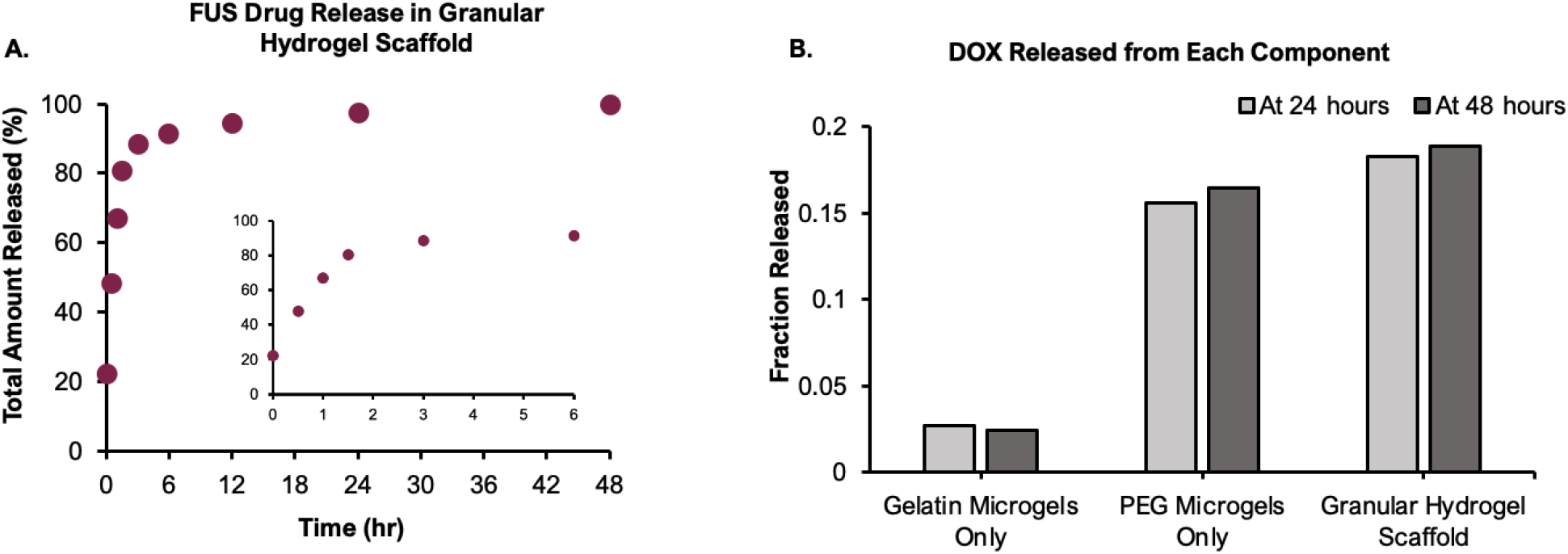
Doxorubucin release within the granular hydrogel scaffold post FUS sonication after 48 hours. **A**. Data was normalized to the amount of drug released in the system after 48 hours. Post-FUS sonication, an initial burst release was followed by controlled release of Doxorubicin. **B**. Each component is assessed for release with the gelatin particle fraction’s burst release accounting for a smaller fraction of the total material release in 24 and 48 hour tests.

The combined release of gelatin and PEG after 24 hours is less than the released observed over 48 (**Figure 4B**), indicating that sonication by FUS does not deplete PEG microgels of drugs. This platform should offer high tunability for controlled release regimes through drug encapsulation in FUS-sensitive microgels (here, gelatin microgels), FUS-insensitive microgels (here, PEG microgels), and the possibility to include drugs in the fiber components (not explored here) or in additional microgel components, which can be designed to have different release profiles to create designed drug-release regimes^38^.

### 2.5 FUS Crosslinking of Granular Hydrogel in Situ

To demonstrate the potential for minimally invasive delivery and FUS-induced stabilization of a granular hydrogel, we investigated FUS-induced crosslinking of granular hydrogels *in situ* in mice cadavers. Using a custom ultrasound guided FUS system, we targeted FUS energy deposition to the center of the granular material injected subcutaneously, then the harvested the implant for gross analysis. We successfully obtained a crosslinked granular hydrogel formed subcutaneously from a minimally invasive injection followed by FUS sonication. Targeted FUS sonication for 15 minutes yielded a robust material which could be explanted and handled (**Figure 5A**). From shear wave elastography imaging (Philips Epiq), we were able to guide delivery of FUS energy and obtain an estimate of modulus before FUS sonication, during FUS sonication at ∼10 minutes (**Figure 5B)**, and post sonication for comparison to pre-crosslinking values. Images from elastography (**Figure 5B**) also allowed real time detection of the focal point of the FUS system and demonstrate the potential for guiding FUS application in larger samples or *in vivo*. Importantly, elastography imaging of the FUS application also provided insight into the crosslinking process through mechanical readouts, providing potential for monitoring and adjusting the FUS process in real time to further reduce the potential for off-target damage. Elastography measurements indicated the granular hydrogel exhibited an elastic modulus of ∼30 kPa before FUS that increased stiffness after FUS treatment to ∼80 kPa (**Figure 5C**). These values correspond in order of magnitude to elastic moduli measured directly and indicate the ability to quantify stiffening during FUS crosslinking. These results confirm our hypothesis that FUS-induced crosslinking, here observed through increases in bulk mechanics of granular hydrogel, can be achieved *in situ* in an implant setting.

**Figure 5.**
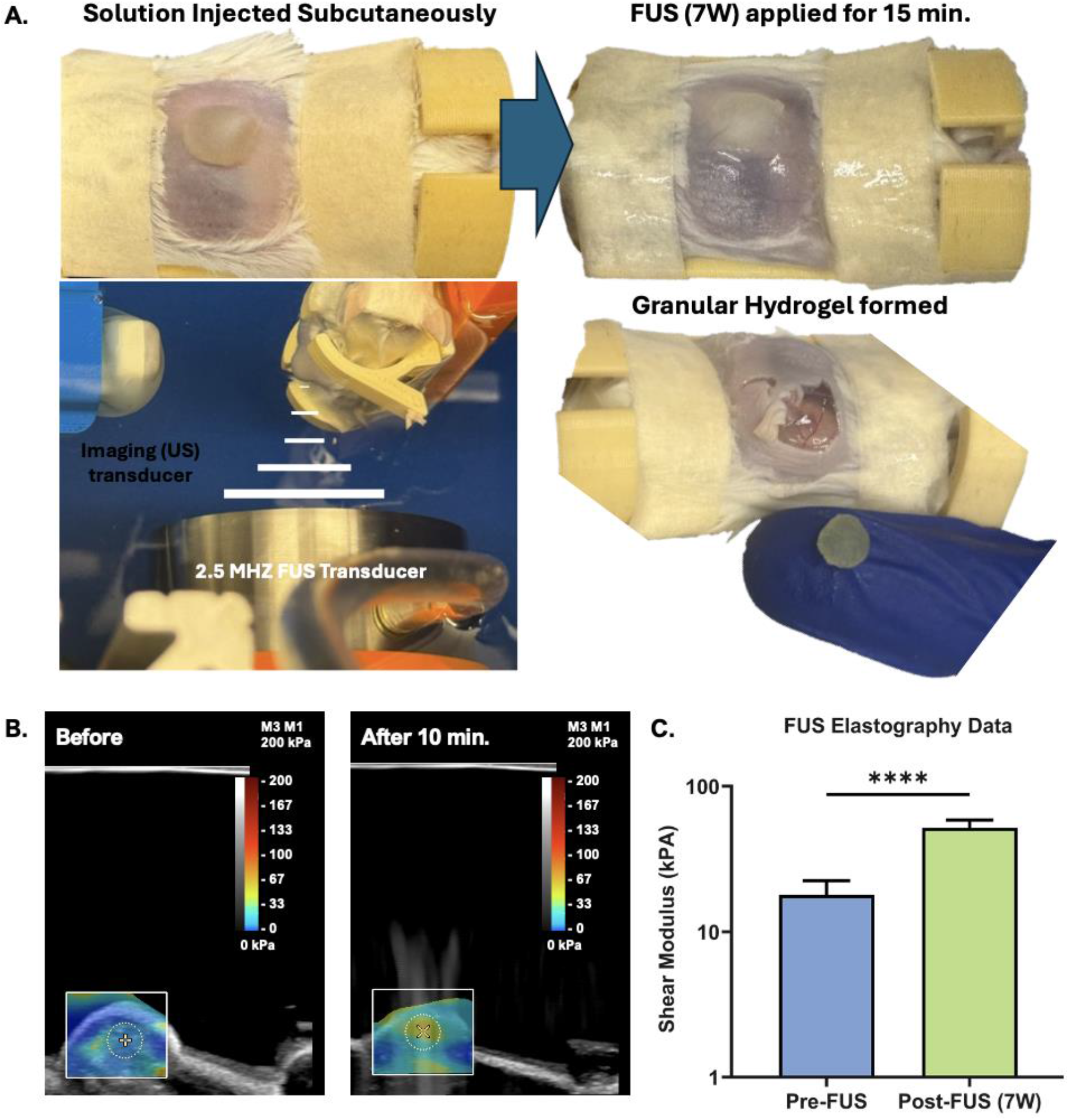
Material FUS crosslinking of the granular hydrogel system *in situ*. **A**. Images of the injected granular hydrogel pre-and post-FUS stimuli. A crosslinked, solid granular hydrogels was obtained post FUS sonications. **B**. Shear Wave elastography ultrasound imaging before (left) and during (right) FUS crosslinking of injected granular hydrogel system **C**. FUS elastography data indicate granular hydrogel stiffness is higher post-FUS compared to prior to FUS.

## 3. Conclusion

Here, we have established a hydrogel biomaterial that can be delivered and stabilized in situ by minimally invasive techniques. Our granular hydrogel system is injectable and porous. Formulation with gelatin microgels allows porosity to be defined in the final scaffold and not be lost during the injection, and the fiber component is crucial to stabilizing and maintain porosity within these materials. Parameters for FUS application were defined to leverage the FUS-responsive chemistry in the granular formulation. The resulting material contains a network of mesoscale pores among microgel and fiber components crosslinked to one another at surfaces. Our results demonstrate that FUS-induced hydrogels exhibit viscoelastic properties comparable to those produced using traditional photo-initiated crosslinking methods. Importantly, we achieved consistent crosslinking across varying FUS power settings, highlighting the robustness and adaptability of this approach. Beyond immediate stability upon injection at the delivery site and porosity, an additional strength of using FUS-crosslinked granular hydrogels as opposed to continuous hydrogels is the potential for engineering design of microgel combinations to support desired biomedical outcomes, as for example in drug delivery applications.

While these advancements are promising, the exact mechanism by which FUS interacts with KPS and induces ROS to facilitate crosslinking remains to be explored. Gaining insight into this mechanism will be essential for optimizing FUS-induced hydrogel formation and ensuring precise control over ROS-mediated reactions, particularly in complex biological environments. While our system demonstrated compatibility with cells introduced to the hydrogel after crosslinking, the effects of FUS stimuli on encapsulated cells, including potential ROS-induced cytotoxicity and cell behavior within the scaffold, remain unexplored. Long-term studies to evaluate biodegradability, mechanical integrity, and overall stability of FUS-crosslinked hydrogels *in vivo* are essential to confirm their suitability for regenerative medicine and wound healing applications.

Despite these limitations, this work lays a strong foundation for the development of FUS-responsive granular hydrogels as a minimally invasive, customizable platform with broad therapeutic potential. By addressing these challenges, future research can enhance this technology, opening new avenues for biomaterial design and focused ultrasound applications in translational medicine.

## 4. Experimental Section

### PEG Microgels and Electrospinning Parameters

Experimental procedures for PEG microgels and electrospun scaffold formation are provided in the Supporting Information.

### Granular Hydrogel Scaffold Fabrication

To create granular hydrogel scaffolds, both PEG microgels and gelatin microgels were first suspended in PBS at 1:1 volume ratio. PEG fibers were suspended in PBS at 1:9 volume ratio. The 300 µL PEG microgels, 100 µL gelatin microgels, and 5% (v/v) PEG fibers were mixed together. 1.5% (w/v) potassium persulfate (KPS, Sigma), 10% (w/v) PEG-thiol, and 2.27% (w/v) PDEGMA were added to the mixture to achieve a final concentration of 0.2%, 0.7%, and 0.16% respectively. This mixture was then centrifuged at 21,130 rcf for 5 minutes and excess solution was aspirated off. To crosslink via focused ultrasound, the material was placed in a water bath under FUS sonication for 15 minutes. To photocrosslink, 10 mM lithium phenyl-2,4,6-trimethylbenzoylphosphinate (LAP) was added in place of KPS to reach a final concentration of 0.7 mM, then crosslinked under 290-320 nm light for 2 minutes at 10 mW/cm^2^. For porosity quantification of scaffolds, PEG microgels were tagged with thiolated rhodamine-B and PEG fibers were tagged with thiolated 5(6)-carboxyfluorescein amidite (FAM) for visualizing. The scaffold was crosslinked, then PBS was added to cover the scaffold, and the scaffolds were cultured in a 37°C, humidified incubator for 20 minutes to liquefy the gelatin microgels prior to imaging.

### Scanning Electron Microscopy (SEM)

The granular hydrogels were flashed frozen with liquid nitrogen then lyophilized overnight. The lyophilized granular hydrogel was placed on a carbon tab in 0.5 inches (12mm) (Al) SEM stubs and imaged on the Quanta 650 (FEI) with a field emission gun source operating in low vacuum mode.

### Focused Ultrasound Machine

FUS was applied to microcentrifuge tubes with a single point sonication using a 1.1-MHz FUS transducer (H107, Sonic Concepts, Bethel Washington), driven by an arbitrary function generator (Tektronix, AFG3052C) and amplifier (E&I, 1040L) with either continuous energy or with 10 millisecond burst lengths and varied pulse repetition rates to result in duty cycles ranging from 25-75% for up to 15 minutes. Both the imaging and treatment transducers were ultrasonically coupled to either the tube containing the precursor solutions or the animal for *in situ* crosslinking using degassed, deionized water at 37°C during the duration of each treatment.

### Treatment of Mice Cadavers with Focused Ultrasound

Mice cadavers were obtained from other studies, shortly after, they were humanely euthanized with ketamine–xylazine according to approved animal care and use protocols for those planned studies, and prior to planned disposal. The subcutaneous space to which our injection was targeted was unaffected by the prior studies. The back of each mouse cadaver was shaved with a hair trimmer and hair remover cream. Cadavers were injected with 0.3 mL of granular hydrogel components (PEG microgels, gelatin microgels, PEG fibers, PEG-thiol solution, KPS, and PDEGMA) via subcutaneous injection with an 18G needle. Mouse cadavers were immediately positioned in a custom ultrasound guided FUS system with a 2.5Mhz transducer (H147, Sonic Concepts). The injected hydrogel precursor was located with the imaging transducer and FUS was then applied to the center of the site with a single point sonication at a 0.6 MPa peak negative pressure for 15 minutes with continuous waveform.

### Rheological Analysis of Granular Hydrogel Properties

Granular hydrogel properties were characterized using rheological studies with a stress-controlled DHR-2 rheometer (TA Instruments), equipped with a 20 mm sandblasted parallel plate. Granular hydrogels were placed onto the bottom plate of the rheometer. Oscillatory tests were performed once the storage modulus reached an equilibrium value.

### Compressive Analysis of Granular Hydrogel Mechanics

Continuous, bulk hydrogel and granular hydrogel properties were also characterized on an Instron using compression at 0.5 mm/second rate with a 10 N load cell on top. The region of 0.1-0.2 strain of each measurement was taken and the stress-strain curve was plotted to find the best fit line, or compressive moduli for the data.

### Statistical Analyses

All results reported with error bars are means with standard deviation. The “*n*” values per group are made by individual data points shown. Statistical significance was assessed at p < 0.05 for all experiments and were calculated using GraphPad Prism. Statistical tests are provided in the figure legends.

## 5. Supporting Information

### Synthesis of poly(di(ethylene glycol) methyl ether methacrylate) (PDEGMA)

Di(ethylene glycol) methyl ether methacrylate (DEGMA, stabilized with monomethyl ether hydroquinone), 4-Cyano-4-(phenylcarbonothioylthio)pentanoic acid (CTA), 4-4’-Azobis(4-cyanovaleric acid) (ACVA, ≥98.0%), and tetrahydrofuran (THF, ≥99.0%, stabilized with butylated hydroxytoluene) were purchased from Sigma-Aldrich. DEGMA was purified by passing the monomer through a column packed with aluminum oxide to remove inhibitor prior to polymerization. All other reagents were used as received. Poly(di(ethylene glycol) methyl ether methacrylate) was synthesized via reversible-addition fragmentation-chain transfer (RAFT) polymerization. DEGMA (2.8 mL, 15 mmol), CTA (55.9 mg, 0.2 mmol), ACVA (5.6 mg, 0.02 mmol) and tetrahydrofuran (8.3 mL, 25 v/v % of monomer concentration) were added synchronously to a 20 mL scintillation vial. The solution was degassed with nitrogen for 30 min, then added to a silicone oil bath set to 70ºC for 4 h. The solution was purified via dialysis against 40/60 methanol/water, then against water. The polymer was then lyophilized and characterized using 1H nuclear magnetic resonance (NMR) spectroscopy and size exclusion chromatography (SEC).

### Electrospun PEG Nanofibers Fabrication

PEG-norbornene electrospun hydrogel nanofibers were produced and adapted from previous literature^28^. Electrospinning hydrogel solution containing 10% (w/v) 8-arm PEG-Norbornene, 5% (w/v) 900 kDa PEO, 4-arm PEG-thiol ([thiol] : [norbornene] = 0.6), and 0.05% Irgacure 2959 in deionized water was dissolved overnight. The 0.6 ratio of thiol to norbornene was chosen to allow for excess norbornenes upon further conjugation of peptides to the fiber surface and secondary crosslinking within the FEMP scaffold. A co-spin solution containing 4% (w/v) 900 kDa PEO in deionized water was dissolved overnight. The hydrogel solution was extruded using a syringe pump at a flow rate of 0.5 mL/hr through a 16G needle to separate individual PEG fibers. The co-spin solution was also extruded using a syringe pump at a flow rate of 0.5 mL/hr through a 16G needle. The second syringe pump was positioned on the opposite side of the mandrel from the hydrogel solution to minimize potentially disruptive interactions between fibers produced from the two different solutions during processing. The fibers were collected on an aluminum foil substrate attached to a mandrel spinning at ∼1,000 rpm. 12 kV positive voltage was applied to the hydrogel needle, 6 kV positive voltage was applied to the co-spin needle, and 4kV negative voltage was applied to the collection substrate.

After collection, the collected fibers on the aluminum foil were crosslinked under UV light for 15 minutes at 5 mW/cm^2^ under nitrogen. Once crosslinked, the fibers were hydrated with deionized water to detach them from the collection substrate. The resulting material was then suspended in 1X PBS. The suspension was homogenized at 10,000 rpm for 2 minutes to segment fibers and then filtered through a 40 μm cell strainer (Fisher) to eliminate aggregations of fibers.

### PEG Microgel Fabrication

Microgels were produced as previously described^39^. Precursor solution containing 6% (w/v) PEG-acrylate (4-arm, 10kDa, JenKem) and 4% (w/v) PEG-thiol (4-arm, 10 kDa, JenKem) in 1X Dulbecco’s phosphate buffered saline (DPBS) was vortexed to dissolve. The solution was placed on a syringe pump at a flow rate of 40 μL/min through a 30G needle into a spinning oil/base solution. Oil/base solution containing light mineral oil, 1% (v/v) Span80, and 0.5% (v/v) triethanolamine (TEOA) spun on a homogenizer at 2,000 rpm. The microgels were left spinning for 1 hour post solution dispensed. Microgels were washed three times with isopropanol via centrifugation at 3,200 rcf for 5 minutes each, then hydrated overnight in 1X DPBS to fully swell. The suspension of microgels were centrifuged at 3,200 rcf for 5 minutes and rehydrated again in 1X DPBS before aliquoting into a 1:1 microgel:1X DPBS ratio.

### Gelatin Microgel Fabrication

Gelatin from bovine skin was dissolved in 1X PBS solution at 15% (w/v) at ∼80°C. The solution was then added to light mineral oil with 2% (v/v) Span 80 heated to ∼80°C and emulsified at 2,000 rpm for 3 minutes. The emulsion was then allowed to cool to room temperature to form gelatin microgels. Once cooled, the gelatin microgels were first centrifuged at 3,000 rcf for 5 minutes. After discarding the supernatant oil, the microgels were suspended in 2-3 mL of 2% (v/v) Pluronic F-127 in 1X PBS to the pellet and centrifuged at 3,200 rcf for 2 minutes. Then, the microgels washed 8x in 1X PBS and centrifuged at 3,000 rcf for 2 minutes after each wash. Finally, the gelatin microgels were filtered through a cell strainer (pluriStrainer, Pluriselect) with mesh sizes ∼1.5X the largest diameter microgel in the distribution (40 µm) to eliminate clusters formed through aggregation.

### Drug Encapsulation in Granular Hydrogel

Doxorubicin (DOX) was encapsulated into gelatin and PEG precursor solutions separately to reach a final concentration of 1 mg/mL. To incorporate DOX into the gelatin microgels, gelatin from bovine skin was dissolved in 1 mg/mL DOX solution at 15% (w/v) at ∼80°C. Gelatin microgels were washed and processed as normal as described in the supplementary information. To incorporate DOX into the PEG microgels, 12% (w/v) 4-arm PEG-acrylate and 8% (w/v) 4-arm PEG-thiol solutions were mixed with 2 mg/mL DOX solution to obtain final concentrations of 6% PEG-acrylate, 4% PEG-thiol, and 1 mg/mL DOX. The PEG microgels were crosslinked and washed. PEG microgels, PEG fibers, and gelatin microgels were placed in microcentrifuge tubes at the desired volume ratios and mixed with PDEGMA, KPS, and PEG-thiol solution as described above. The granular hydrogel material underwent FUS sonications for 10 minutes. Immediately, 1X PBS was placed on the samples to obtain a T0 time point for DOX released. Controlled volumes of PBS were pipetted out of the tube at various timepoints over 48 hours.

### Quantification of Drug Release

Samples taken from each timepoint of the drug release experiment were saved in microcentrifuge tubes and placed in the fridge until quantification. All samples were pipetted into a 96 wellplate for absorbance measurements. These measurements were conducted on plate reader (Multimode Microplate, Tecan Spark) at 475 nm light to calculate the absorbance values of DOX within each sample. A standard curve was generated by preparing DOX solutions of known concentrations as low as 0.006 µg/mL. A best fit curve was obtained from plotting absorbance values of the known concentrations in GraphPad Prism with an r-squared value of 0.998. The best fit equation was used in DOX release experiments to obtain an amount of material in solution at each time point. Further calculation was conducted to get the percent fraction of DOX released over time within the system.

### Continuous Hydrogel Fabrication

A continuous, bulk hydrogel was created using a 20% (v/v) PEG-diacrylate (PEGDA) solution (700 Da, Sigma), 2.27% (w/v) PDEGMA, and 1.5% (w/v) KPS solutions. This material was placed under FUS sonication until a hydrogel formed. To photocrosslink the continuous hydrogel, 10 mM LAP was added in place of KPS as described above, then crosslinked for 2 minutes under 290-320 nm at 10 mW/cm^2^.

### Image Analysis and Porosity Quantification

Images were taken using a Leica DMi8 widefield microscope and Leica Stellaris 5 confocal microscope. Porosity images were processed using FIJI software.

### Cell Staining

In experiments measuring cell viability, cells were stained using a LIVE/DEAD assay kit (ThermoFisher, L3224) that used calcein-AM and ethidium homoimer-1 to stain live and dead cells, respectively.

**Supporting Video 1**. Packed gelatin microgels pre-FUS

**Supporting Video 2**. Melted gelatin microgels post-FUS

**Supporting Figure 1**. Schematic of RAFT synthesis of PDEGMA

**Supporting Figure 2**. ^1^H NMR (400 Mhz, DMSO-d6) spectrum of PDEGMA

**Supporting Figure 3**. Chromatogram of PDEGMA obtained by RAFT polymerization

**Supporting Table 1**. Characterization information for PDEGMA

## Acknowledgements

Funding from NIH NIGMS R35GM147410 and NIH NIGMS T32GM136615.

## Notes

### Competing Interest Statement

The authors have declared no competing interest.

