## Supporting Information for "Focused Ultrasound Crosslinkable Granular Hydrogels"

Natasha L. Claxton[1], Estelle He[3], Kelly Bukovic[2], Eric A. Thim[1], Rachel A. Letteri[2], Steven R. Caliari[1,2], Richard J. Price[1], Matthew R. DeWitt[1 *], & Christopher B. Highley[1,2 *]

[1]Department of Biomedical Engineering, University of Virginia, Charlottesville, VA 22903, USA
[2]Department of Chemical Engineering, University of Virginia, Charlottesville, VA 22903, USA
[3]Department of Physics, University of Virginia, Charlottesville, VA 22903, USA

*
*

**Supporting Video 1.** Packed gelatin microgels containing black dye prior to FUS treatment exhibiting solid-like behavior.

**Supporting Video 2.** Packed gelatin microgels containing black dye after FUS treatment, where they have melted and are flowing as a liquid.

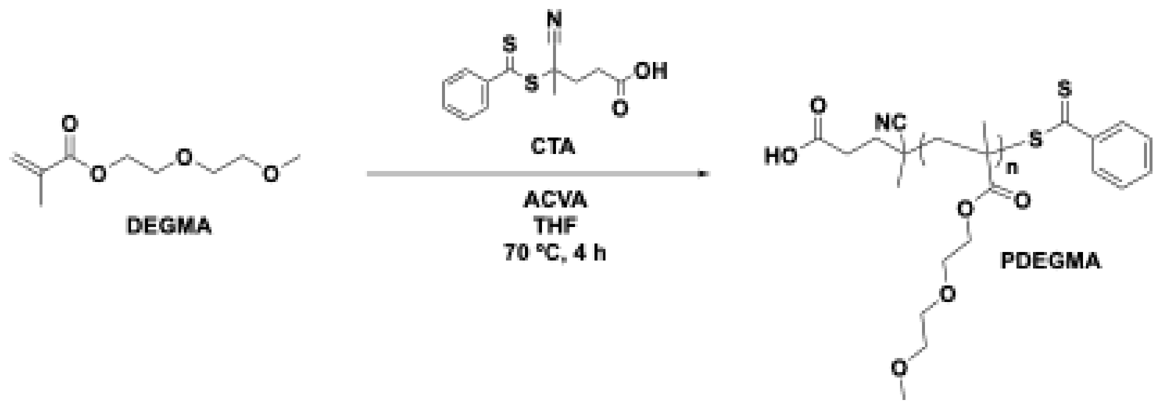


**Supporting Figure 1.** Schematic illustrating RAFT synthesis of PDEGMA.

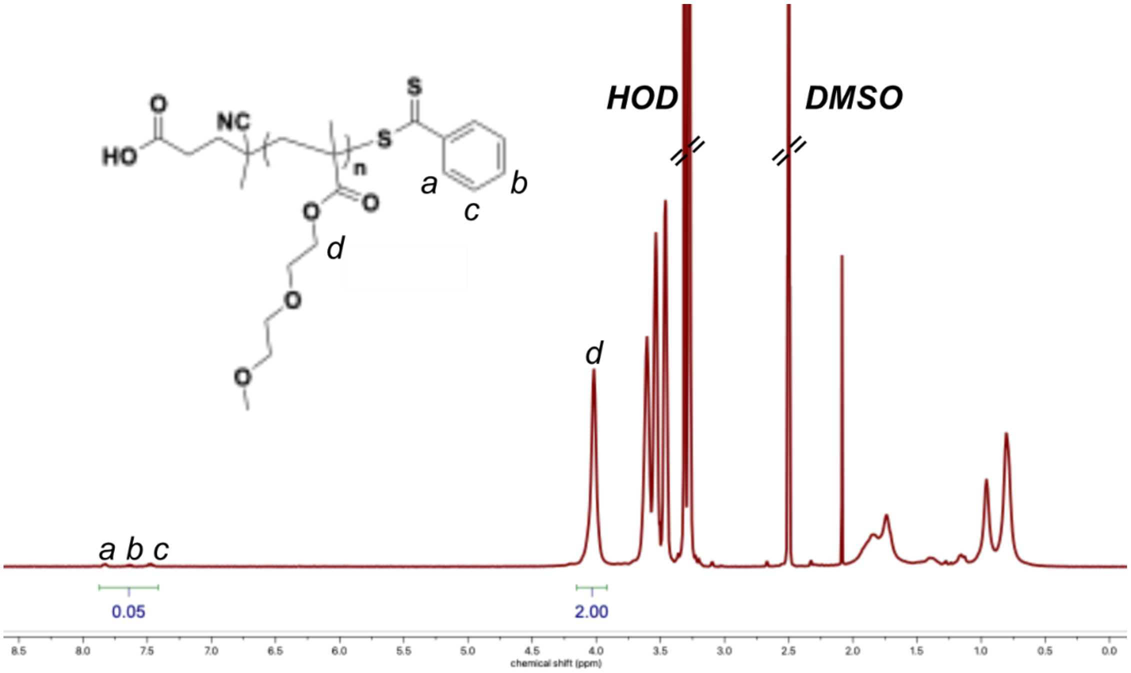


**Supporting Figure 2.** [1]H NMR (400 Mhz, DMSO-d6) spectrum showing characterization of PDEGMA to determine molecular weight by comparing integrations of the CTA endgroup (peaks a, b, and c) to DEGMA repeats (d) (see Supporting Table 1).

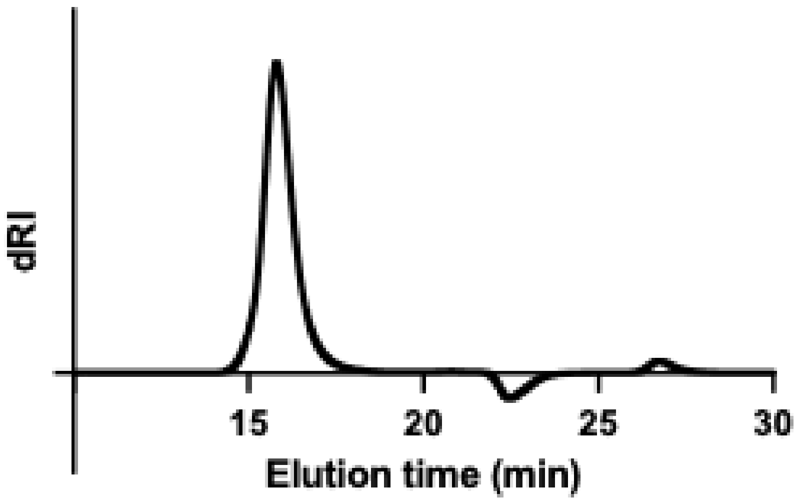


**Supporting Figure 3.** Chromatogram of PDEGMA obtained by RAFT polymerization, used to quantify molecular weight against PMMA standards (see Supporting Table 1).

| $M_{n, NMR}$ (kDa) [a] | $M_{n, SEC}$ (kDa) [b] | Đ [b] |
| --- | --- | --- |
| 19.1 | 20.8 | 1.28 |

**a** Determined by $^1H$ NMR spectroscopy from the relative integrations of the CTA end group peaks at 7.4-7.9 ppm and of DEGMA repeat units at 4.0 ppm. **b** Estimated by SEC in TFE with 0.02 M NaTFAc relative to PMMA standards.

**Supporting Table 1.** Characterization of molecular and polydispersity index of PDEGMA from NMR and size exclusion chromatography.
